# *In ovo* electroporation of chicken limb bud ectoderm

**DOI:** 10.1101/2020.11.18.388033

**Authors:** Reiko R. Tomizawa, Clifford J. Tabin, Yuji Atsuta

**Affiliations:** Department of Genetics, Harvard Medical School, Boston, MA, USA; Department of Biology, Faculty of Sciences, Kyushu University, Fukuoka, Japan

**Keywords:** Limb bud ectoderm, *In ovo* electroporation, AER, Time-lapse imaging, Wound healing, Rac1 GTPase

## Abstract

Deciphering how ectodermal tissues form, and how they maintain their integrity, is crucial for understanding epidermal development and pathogenesis. However, lack of simple and rapid gene manipulation techniques limits genetic studies to elucidate mechanisms underlying these events. Here we describe have an easy method for electroporation of chick embryo limb bud ectoderm, enabling gene manipulation during ectoderm development and wound healing. Taking advantage of a small parafilm well that constrains DNA plasmids locally and the fact that the limb ectoderm arises from a defined site, we target the limb ectoderm forming region by *in ovo* electroporation. This approach results in efficient transgenesis of the limb ectodermal cells. Further, using a previously described *Msx2* promoter, gene manipulation can be specifically targeted to the apical ectodermal ridge (AER), a signaling center regulating limb development. Using the electroporation technique to deliver a fluorescent marker into the embryonic limb ectoderm, we show its utility in performing time-lapse imaging during wound healing. This analysis revealed previously unrecognized dynamic remodeling of the actin cytoskeleton and lamellipodia formation at the edges of the wound. We find that the lamellipodia formation requires activity of Rac1 GTPase, suggesting its necessity for wound closure. Our method is simple and cheap, and permits high throughput tests for gene function during limb ectodermal development and wound healing.

## 1 INTRODUCTION

Embryonic epithelial tissues have pivotal roles in organizing tissue development as well as in physically maintaining organ shape. Because of its high accessibility, the chick limb bud ectoderm has served as a model to study how ectodermal cells interact with the underlying mesoderm to coordinate organogenesis, and in investigation of how epidermal tissue heals following a mechanical wound (Fernandez-Teran and Ros, 2008; Fernandez-Teran et al., 2013; Martin and Lewis, 1992; Brock et al., 1996). For gene delivery to the chick limb ectoderm, an RCAS vector, a replication-competent avian retrovirus, is commonly used (Hughes et al., 1987; Logan et al., 1997; Kengaku et al., 1998). Although infection of RCAS yields high efficiency of expression of exogenous genes, its use has several limitations. First, gene expression from RCAS requires time to commence (typically 2-3 days), thus it is unsuitable for examining earliest events of limb bud formation including initiation of the apical ectodermal ridge (AER) development, which is essential for proximodistal patterning of the limb bud. Secondly, due to its replication capability, the viral particles are readily spread outside of targeted tissues, thereby complicating interpretation of resultant phenotypes (Fekete and Cepko, 1993). Lastly, the packaging capacity of RCAS vector is relatively limited, and it is not feasible to accommodate cDNA larger than ~2 kbp. Other alternative options for gene manipulation of the chick limb ectoderm include the use of replication incompetent viruses such as lentiviruses or generation of transgenic chicken lines using a surface ectoderm specific driver (Beronja et al., 2010; Li et al., 2000). Yet, these methods are, themselves, problematic. For instance, lentiviral infection often results in low efficiency and slow expression of introduced genes, while making transgenic chickens is time and resource consuming. Therefore, establishment of an effective, rapid, simple, and cheap gene manipulation technique for the chick limb ectoderm has been desired.

*In ovo* electroporation is widely used for gene manipulation in avian embryonic tissues. This approach allows transfection of both proliferating and non-dividing cells. Moreover, as a transgene is directly introduced in the form of a DNA vector, gene expression is triggered more rapidly than in retroviral infection. However, electroporation often causes severe tissue malformations due to the excessive electric voltage required (Scaal et al., 2004). It can also be difficult to achieve focal gene delivery because the DNA solution applied to the embryo can easily spreads. In the current study, we have devised a new electroporation method for the chick limb ectoderm by taking advantage of a small well made of stacked parafilm. The use of the well not only prevents diffusion of the DNA plasmid solution, it also acts as a voltage barrier mitigating embryonic damage, thereby improving the viability of electroporated embryos while successfully targeting the prospective limb field. We also show that AER specific gene manipulation can be achieved by exploiting a previously described *Msx2* promoter. Blocking AER formation by misexpression of Noggin abrogates *Msx2* activity. The ectoderm-targeting electroporation method described here can also be used to potentiate *in vivo* time lapse analysis of processes such as wound healing. Using this approach to visualized cellular behavior and actin dynamics during ectodermal wound closure, we show that wound ectodermal cells extended lamellipodia to bridge a gap created in the tissue, driven by the activity of the Rac1 small GTPase. Taken together, this parafilm-well electroporation has opened a way to investigate the cellular and molecular mechanisms underlying limb ectoderm development, and the process of epithelial wound healing in chickens.

## 2 RESULTS AND DISCUSSION

### 2.1 Lineage tracing of presumptive limb bud ectoderm in chicken embryos

As a first step towards developing a method for gene-manipulation of limb bud ectodermal cells, we needed to definitively locate the prospective limb ectoderm prior to limb initiation. We focused on HH10 embryos since electroporation at this stage would allow sufficient time for gene expression before AER formation, marked by initiation of *Fgf8* expression at HH16 (Fernandez-Teran and Ros, 2008). A previous study mapping the origins of the limb bud mesenchyme showed that lateral plate mesoderm adjacent to Hensen’s node in HH10 embryos gave rise to the limb and flank mesenchyme at HH20 (Matsubara et al., 2017), thus we targeted similar regions of the same staged embryos (Fig. 1A and B). PKH26, a red fluorescent membrane dye, was used to label small regions of surface ectodermal cells that were located about 300 μm laterally from the posterior neural tube. As a result, we found that regions of surface ectoderm located approximately 600, 900, and 1300 μm posterior to the last somite at HH10 gave rise to the forelimb, flank and hindlimb ectoderm cells at HH20, respectively (Fig. 1A and B), consistent with previous reports but providing more precise resolution (Altabef et al., 1997).

**Figure 1.**
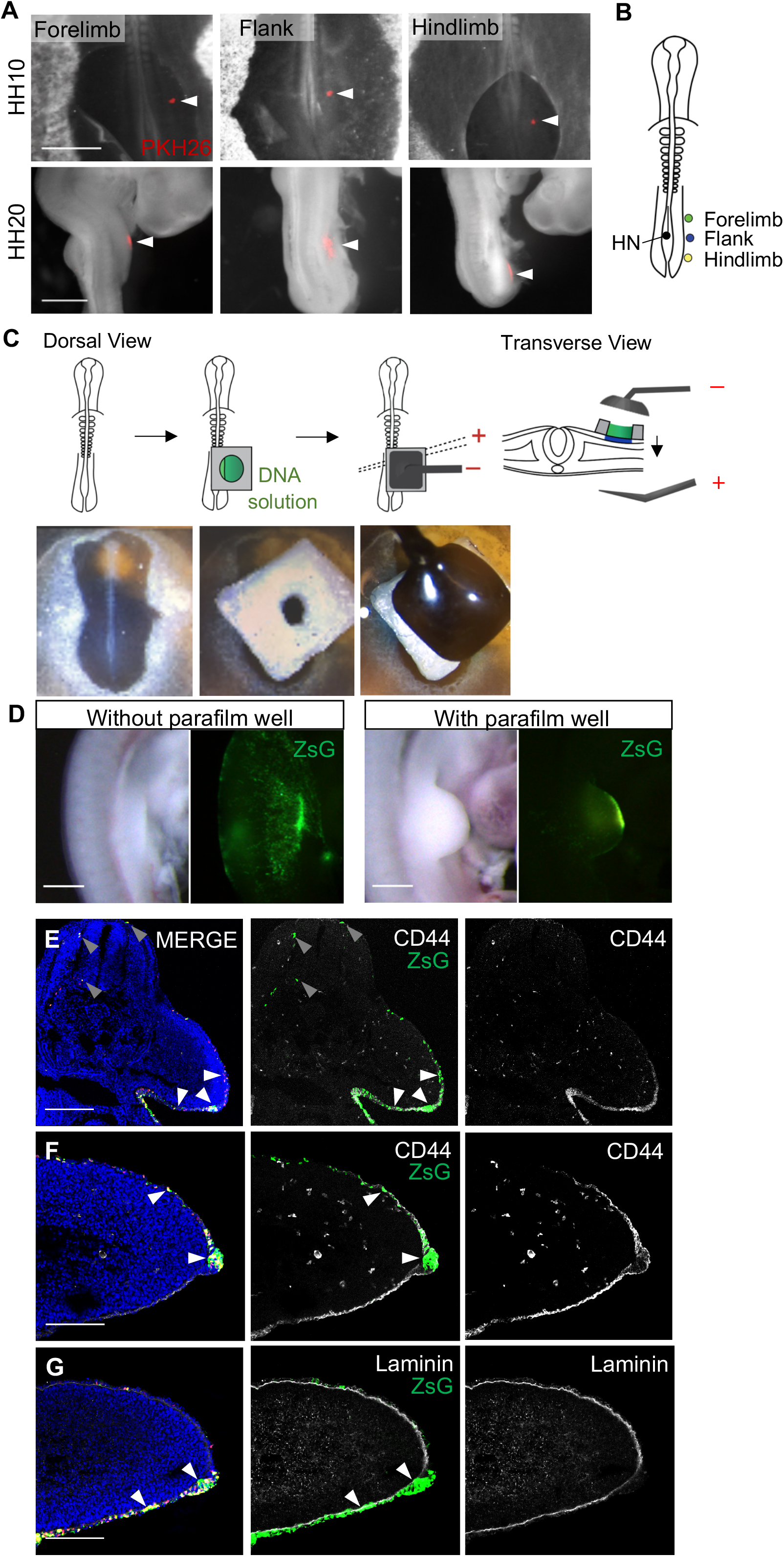
Transgenesis of chicken limb bud ectoderm by *in ovo* electroporation. (A) The surface ectodermal cells in HH10 embryos were stained with PKH26 (a red fluorescent dye) marked with white arrowheads (upper panels). Stained regions at HH20 are indicated with white arrowheads (lower panels). (B) A schematic of results of the lineage tracing (n = 3 for each). HN stands for Hensen’s node. (C) Upper panels are schematics showing use of a parafilm well to target prospective forelimb ectoderm in HH10 embryos. The parafilm well was placed on an embryo, subsequently the well was filled with plasmid DNA solution, and then electric pulses were applied. Lower panels are actual views. (D) Electroporation of a plasmid expressing H2BmCherry-ires2-ZsGreen1 (H2BmCh-ZsG) in the forelimb ectoderm either without or with the parafilm well was performed. Note that only ZsG signals are shown. (E) Transverse sections of HH20 embryos electroporated with H2BmCh-ZsG using the parafilm well, and stained with antibodies against CD44. White arrowheads indicate electroporated AER cells while gray ones point to ZsG signals in the neural ectoderm. (F, G) Transverse sections of electroporated HH20 forelimbs stained with antibodies for CD44 (F) and Laminin-1 (G). White arrowheads mark ZsG signals in the limb ectoderm that are colocalized with the markers. Scale bars: 1 mm in (A, D), 400 μm in (E), 200 μm in (F, G).

### 2.2 Optimization of conditions for electroporation into chicken limb ectodermal cells

Next, we optimized electroporation into the prospective limb ectoderm of HH10 embryos. Various electroporation parameters were tested to determine the optimal conditions for inducing gene expression, utilizing a ZsGreen1 (ZsG) reporter, in limb ectodermal cells at HH20 (Table 1). Electroporation, after injecting the DNA plasmid solution beneath the vitelline membrane, resulted in malformed limb buds; consistent with previous reports that electric voltage can cause considerable stress to the embryos (Fig. 1C, D and Table 1) (Scaal et al., 2004), which in this case may have been exacerbated by the close proximity of the negative electrode to the embryos. Additionally, we saw widespread signal rather than localized ectodermal expression, possibly caused by the spread of the DNA solution to other tissues (Fig. 1D). To mitigate these issues, we made a small parafilm well with a thickness of ~1 mm, which created a seal around the ectoderm to prevent DNA leakage. This also had the effect of increasing the distance between the electrode and the embryo tissue (Fig. 1C). Parafilm was selected over other materials, such as filter paper, due to its water repellent properties, and the fact that it could insulate against electrical damage. Using the parafilm well, indeed resulted in lower percentage of malformed embryos (12~30 % improvement) and higher localization of ZsG signals (Table 1, Fig. 1D). Based on the percentage of viable embryos (number of samples that did not show malformations out of total embryos electroporated) and efficiency (embryos with signal in the limb ectoderm out of total embryos electroporated), we determined that three poring pulses at 30 V for 0.1 msec was optimal for this experiment. To confirm that expression in embryos electroporated with the parafilm well was confined to the ectodermal compartment, we sectioned and stained these samples with CD44 and Laminin, which marks the limb ectoderm and basal lamina, respectively (Fig. 1E-G). The ZsG signal was seen throughout the limb ectoderm, including the AER, while it was absent from the mesenchyme (Fig. 1E-G). Additionally, some signal was detected in ectodermal derivatives outside of the limb field, such as in the neural tube (Fig. 1E).

**Table 1.** Optimization for electroporation parameters. Fluorescent signals were checked when embryos grew to stage HH20. All conditions used 8 transfer pulses of 5 V, 10 msec. Embryos with signals in the limb bud were categorized under positive signals.

| Poring Voltage (V) | Pulse Length [msec] | No. of Pulses | Parafilm Well | Total | Malformed | % Viability | Pos. Signals | % Efficiency |
| --- | --- | --- | --- | --- | --- | --- | --- | --- |
| 20 | 0.2 | 3 | Yes | 26 | 14 | 46 | 15 | 58 |
|  |  |  | No | 25 | 20 | 20 | 19 | 76 |
| 30 | 0.1 | 3 | Yes | 25 | 17 | 32 | 17 | 68 |
|  |  |  | No | 25 | 20 | 20 | 20 | 80 |
| 40 | 0.1 | 3 | Yes | 26 | 16 | 39 | 12 | 46 |
|  |  |  | No | 23 | 21 | 9 | 15 | 65 |

### 2.3 A murine AER-specific Msx2 promoter drives gene expression in chick AER

Next, we tried to specifically target exogenous gene expression to the AER. The AER plays pivotal roles in limb outgrowth and patterning by secreting morphogens such as Fgfs and Wnts (Fallon et al., 1994; Crossley et al., 1996; Kawakami et al., 2001). Our electroporation method induces gene expression throughout the limb ectoderm. To localize transgenic activity specifically to the AER, we decided to take advantage of a previously described 439 base pair AER-specific promoter, derived from the mouse *Msx2* gene (Liu et al., 1994). To see if the element would also drive expression in the chicken AER, the mouse *Msx2* promoter (hereafter *Msx2*) was inserted upstream of human placental alkaline phosphatase (PLAP) (Billings et al., 2010). In the control embryos with only a minimal TATA-box, alkaline phosphatase staining was absent (Fig. 2A). In contrast, in *Msx2*-PLAP electroporated samples, cells the AER were positive for alkaline phosphatase staining. Moreover, this staining was confined to the AER, while many cells outside of the AER expressed mCherry driven from an electroporation-control vector, suggesting that *Msx2* activity is specific to the AER in chickens as shown in mice (Fig. 2B). We further checked the AER specificity of the *Msx2* promoter by looking at transverse sections of embryos electroporated with a plasmid carrying *Msx2*-DsRed along with the ZsG control vector. ZsG signals can be seen throughout the limb ectoderm and in the neural ectoderm (Fig. 2C-E upper panels). However, DsRed signals driven by *Msx2* were exclusively found in the AER, colocalized with endogenous Msx protein (Fig. 2C-E; lower panels). DsRed was also occasionally observed in the ventral limb ectoderm (Fig. 2E). A previous study using the same *Msx2* promoter showed a similar expression pattern of *Msx2* activity in the AER and ventral ectoderm in mice. The authors suggested that the AER ventral boundary in mice is not specified until E10.5 (Kimmel et al., 2000). Similarly, our results imply that the ventral boundary of the AER in chick may not be specified until after stage HH20. The *Msx2*-reporter construct was also introduced into the limb mesenchyme of HH20 embryos, which express Msx2 in its distal and antero-posterior portions. (Fernández-Terán et al., 2006; Fig. 2F). However, the *Msx2* promoter did not drive DsRed expression in the limb mesenchyme, confirming that *Msx2* activity is specific to AER amongst Msx2 expressing tissues, at least, in the limb fields (Fig. 2F).

**Figure 2.**
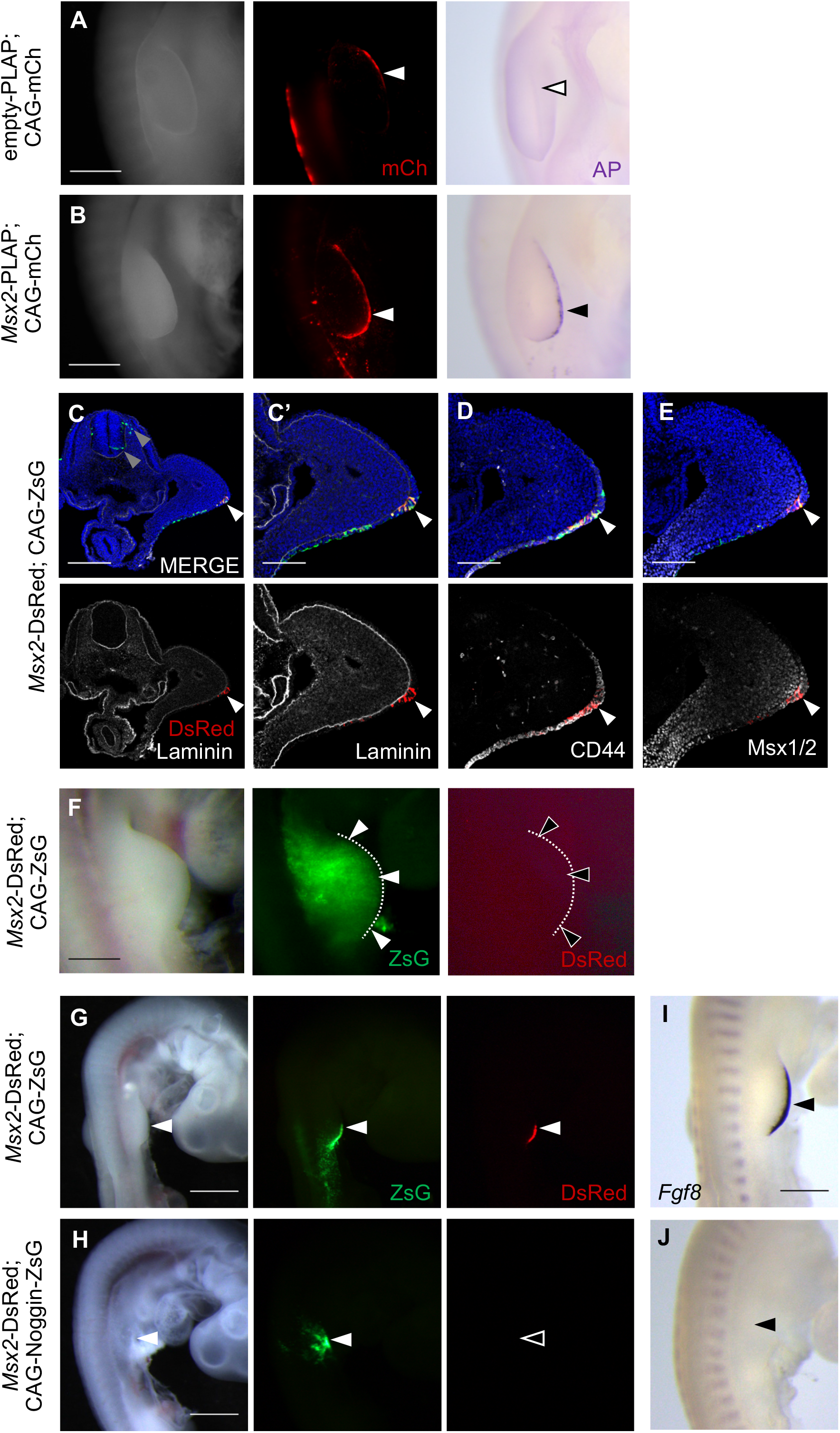
Chicken AER-specific gene manipulation by using mouse *Msx2* promoter. (A) Electroporation of the plasmid carrying a minimal promoter/placental alkaline phosphatase (empty-PLAP) or a *Msx2* promoter/PLAP (*Msx2*-PLAP) into chick forelimb ectoderm along with a mCherry (mCh) expressing plasmid, followed by AP staining. A black arrowhead in (B) indicates positive AP staining signals at the AER. (C-E) Sectioned limbs electroporated with *Msx2*-DsRed and CAGGS-ZsGreen1 (CAG-ZsG) vectors, and stained with antibodies against Laminin1 (C, C’), CD44 (D), and Msx1/2 (E). Gray arrowheads show ZsG signals in the neural tube, and white arrowheads mark DsRed signals in the AER. (F) Co-electroporation of *Msx2*-DsRed and CAG-ZsG into the limb mesenchyme. *Msx2*-DsRed yielded no signals, indicated by blank arrowheads. (G-J) Overexpression of Noggin cDNA in the right forelimb ectoderm disrupts forelimb formation, which is represented by losses of *Msx2* promoter activity and *Fgf8* expression. (G, H) The control (CAG-ZsG; F) or Noggin plasmid (CAG-Noggin-ZsG; H) was co-electroporated with *Msx2*-DsRed. Forelimb forming regions are indicated by arrowheads. (I, J) Whole mount *in situ* hybridization for *Fgf8* mRNA in control and noggin electroporated HH20 chick embryos. Black arrowheads mark electroporated regions. Scale bars: 1 mm in (A, B, F, G, H), 500 μm in (I), 400 μm in (C), 200 μm in (C’-E).

The AER forms along the distal dorsal-ventral border of the limb bud. To verify that *Msx2* activity is specific to the functional AER, as opposed to cells at the border location, we tested the ability of *Msx2* to drive expression in limb buds where AER formation is inhibited. to accomplish this, we misexpressed noggin, an antagonist of BMP signaling, which is known to be required for AER formation (Ahn et al., 2001; Pizette et al., 2001). Thus, a noggin-expressing plasmid was co-electroporated with *Msx2-*DsRed (Fig. 2G-J). In controls, in which the noggin construct was not included, DsRed signal is present in the AER, co-localizing with the AER marker *Fgf8* (Fig. 2G, I). However, in noggin-electroporated samples both DsRed fluorescence and *Fgf8* mRNA were absent (Fig. 2H, J), and as expected in the absence of an AER, the forelimb failed to develop properly. These results suggest that *Msx2* promoter activity reflects the status of AER formation, and by combining the electroporation technique described here with the use of the *Msx2* promoter, we are able to manipulate gene expression specifically in cells of the AER.

### 2.4 Time-lapse imaging of chicken limb ectodermal tissue identifies lamellipodia formation, regulated by Rac1, during wound closure

Fluorescent protein-labeling of the limb ectodermal cells enabled us to directly visualize the behavior of individual cells *in vivo*. Using a modified New culture method (Chapman et al., 2001), an HH20 embryo electroporated with a Lifeact-mRuby expressing plasmid, labeling filamentous actin (F-actin; Riedl et al., 2008), was placed in a glass-bottom dish and followed by time lapse confocal microscopy (Fig. 3A). In this instance, the hindlimb bud ectoderm was targeted for electroporation, instead of the forelimb, in order to avoid vibratory influence from the beating of the embryonic heart. At HH20, the limb ectoderm simply consists of two layers; a single basal layer of cuboidal cells and an outer layer of squamous cells called periderm (Martin and Lewis, 1992). Our electroporation technique results in the transfection of both types of cells, because the cells are derived from a single progenitor population. However, we focused on the periderm because they exhibit the typical polygonal epithelial appearance, and their visualization was easier at a single cell level (Fig. 3B). Our imaging system revealed the morphology and ultrastructure of individual cells, including intra- and supracellular actin networks that were dynamically remodeled despite their static shape (Fig. 3B, mov. S1). This observation prompted us to examine cellular behavior during wound closure, monitoring changes in cell shape at high resolution. Of note, we found that leading edge cells protruded actin-rich lamellipodia, which crawled on the limb mesenchyme to bridge the gap caused by the wound (Fig. 3C, 3C’, and mov. S2), in addition to supracellular actin cables that have been previously reported (Martin and Lewis, 1992). The wound healing of embryonic chick limb epidermis has been describes as being achieved through a “purse-string” mechanism driven by an actomyosin contractile cable (Martin and Lewis, 1992; Brock et al., 1996; Begnaud et al., 2016). However, our results imply that the process of chick epidermal wound healing may be more complex, requiring the combinatorial action of an actomyosin purse-string and lamellipodia-based cell crawling, as seen in other contexts of wound healing such as in mouse and human corneal epithelium (Danjo and Gipson, 1998; Klarlund, 2012). One possibile reason that the lamellipodia activity was not previously identified during limb ectoderm healing, is that such fine cellular structures can easily be disrupted by conventional fixation (Sanders et al., 2013).

**Figure 3.**
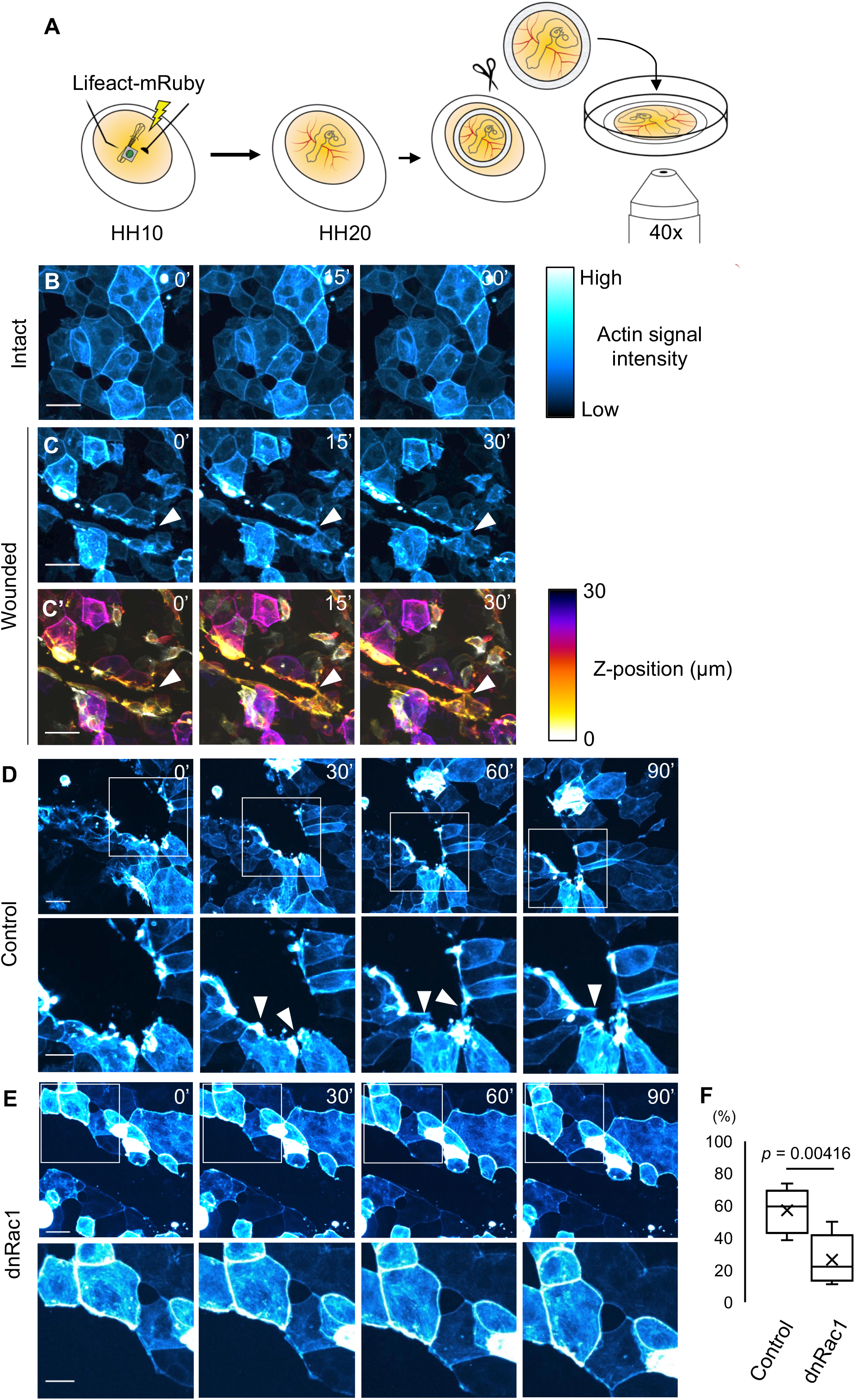
Actin-rich lamellipodia formation during wound healing that requires activity of Rac1. (A) An HH20 chick embryo electroporated with Lifeact-mRuby expressing plasmids was placed in a glass-bottom dish, and subjected to time-lapse imaging analyses by confocal microscopy. (B) Selected frames from the time-lapse movie (Movie S1). F-actin signals were color-coded according to the intensity. (C, C’) Time-lapse imaging show that the wound epithelial cells bridged a lesion by pseudopodia, which is marked by white arrowheads (see also Movie S2). F-actin signals were depth-coded in (C’). Pseudopodia exist at a lower height, implying that these structures are crawling on non-labelled limb mesenchyme. (D-F) Rac1 activity is necessary for lamellipodia formation. (D) Control leading edge cells seal a lesion by extending lamellipodia (white arrowheads). Magnified stills are shown as lower panels. (E) dnRac1 transfected cells hardly display lamellipodia. See also Movies S3 and S4. (F) Quantitative representation of the number of the cells with lamellipodia per the electroporated cells at a wound edge. 63 cells out of 113 control-electroporated cells in 5 embryos showed lamellipodia, while 20 cells per 84 dnRac1 expressing cells in 5 embryos extended lamellipodia during time-lapse imaging. The *P* value was obtained using a 2-tailed, unpaired Student’s *t* test. Scale bars: 25 μm in (B, C, D, E), 15 μm in magnified images of (D, E).

Rac1, a small GTPase, is a major actin cytoskeletal regulator driving lamellipodia activity (Nobes and Hall, 1995), and activation of Rac1 is required for lamellae formation during mouse keratinocyte wound healing (Tscharntke et al., 2007). Therefore, we investigated whether the function of Rac1 is necessary for the lamellipodia formation in chick limb wound epithelia. To that end, we overexpressed a dominant-negative form of Rac1 (dnRac1) in limb ectodermal cells, to inhibit endogenous Rac1 during wound healing (Kuroda et al., 1998). In the control samples, the cells at the wound edge actively extended lamellipodia toward the gap, thereby narrowing and sealing the lesion (Fig. 3D, mov. S3). In contrast, dnRac1 expressing cells exhibited significantly fewer lamellipodia, compared to the control cells, during mechanical wound closure (Fig. 3E, F, mov. S4). Together, by taking advantage of the electroporation technique described herein, we demonstrated that there is an extension of lamellipodia by the leading-edge cells during chick limb epithelial wound healing, which is controlled by the activity of Rac1.

## 3 EXPERIMENTAL PROCEDURE

### 3.1 Experimental animals

White leghorn eggs were obtained from Charles River. Chicken embryos were staged according to Hamburger and Hamilton stages (HH, Hamburger and Hamilton, 1992). All animal experiments were performed under the guidelines of the Harvard Medical School Institutional Animal Care and Use Committee.

### 3.2 PKH26 staining

PKH26 dye (1:10 dilution, Sigma-Aldrich) was injected onto the ectoderm of HH10 chicken embryos (about 38 hours incubation at 38 °C), imaged immediately, and re-incubated for 2 days. At HH20, embryos were dissected and imaged again.

### 3.2 *In ovo* electroporation and expression vectors

Chicken embryos were electroporated at stage HH10. The well was made with 8 layers of parafilm and a 1 mm diameter hole. DNA solution at a concentration of 4.5 μg/μL was injected either below the vitelline membrane above limb ectoderm or into the parafilm well, which was placed directly onto the ectoderm. 3 poring pulses of 20, 30, or 40 V for 0.2 or 0.1 msec followed by 8 transfer pulses of 5V, 10 msec were used (Super Electroporator NEPA21-type II, NEPA GENE). pT2A-CAGGS-H2BmCherry-ires2-ZsGreen1 (ZsG), pT2A-CAGGS-ires2-ZsG, pT2A-CAGGS-Noggin-ires2-ZsG and pT2A-CAGGS-dnRac1-ires2-ZsG were made in (Atsuta and Takahashi, 2016). For pStagia3-*Msx2*, the mouse *Msx2* AER-specific promoter sequences (Liu et al., 1994), which were amplified from mouse genomic DNA, were inserted into pStagia3 (a gift from Dr. Constance L. Cepko [Harvard Medical School]; Billings et al., 2010). To make pSta-*Msx2*-DsRed, two of the *Msx2* promoters were subcloned tandemly into pStagia3, then GFP-ires2-PLAP was replaced with cDNA of DsRed-Express2 (Clontech). To obtain pT2A-CAGGS-Lifeact-mRuby2-ires2-ZsG, Lifeact-mRuby2 CDS was amplified by PCR from mRuby2-Lifeact-7 (addgene #54674; Lam et al., 2012), and subcloned into pT2A-CAGGS-ires2-ZsG. pCAGGS-mCherry was provided by Dr. Constance L. Cepko.

### 3.3 Immunostaining

Immunological staining on histological sections were performed as described in (Atsuta and Takahashi, 2016). HH20 chicken embryos were dissected and fixed in 4% PFA for 3 hours, placed in a series of sucrose/PBS solution, and embedded in OCT compound (Sakura Finetek). 12 μm cryo-sections were collected and permeabilized with 0.5% Triton X-100/PBS for 10 min. Sections were incubated with blocking buffer (1% Blocking reagent [Roche]/TNT) for one hour followed by primary antibodies overnight at 4 °C. On day 2, sections were washed with TNT buffer and incubated with secondary antibodies for 5-6 hours at 4°C. Vectashield mounting media with DAPI (Vector Laboratories) was used before imaging with a confocal microscope (Zeiss LSM710). The following primary antibodies and concentrations were used: mouse anti-CD44 (1:500, Bio-Rad MCA5762), mouse anti-Laminin (1:500, DSHB 3H11), mouse anti-Msx1/2 (1:100, DSHB 4G1) and rabbit anti-RFP (1:500, Abcam ab62341).

### 3.4 Alkaline phosphatase (AP) staining

AP staining with chicken embryos were performed as previously described in (Billings et al., 2010). HH20 embryos were dissected and fixed in 4% PFA for 45 mins at RT followed by incubation at 65 °C for 1.5 hrs. Embryos were rinsed with detection buffer to raise the pH and incubated in developing solution at RT for 1.5 hrs. Coloring reaction was stopped with PBS washes and embryos were post-fixed with 4% PFA for 20 min before imaging.

### 3.5 Whole mount *in situ* hybridization (WISH)

WISH was performed as previously described with slight modifications (Tonegawa et al., 1997). Electroporated HH20 embryos were dissected and fixed in 4% PFA for 3 hours and stored in methanol. On day 1, embryos were re-hydrated in gradual replacement with PBST then permeabilized with Proteinase-K (20mg/ml) for 15 minutes and re-fixed for 20 minutes. After 3 washes with PBST, embryos were pre-hybridized in ULTRAhyb (Invitrogen) at 68 °C for one hour, then were hybridized with RNA probes overnight. On day 2, samples were washed with hybri-wash buffer and MABT, followed by blocking with 10% Blocking reagent /MAB and FBS. Secondary antibodies (Anti-dig-AP, Roche) were added for overnight incubation at 4 °C. On day 3, embryos were washed in MABT buffer for one hour and NTMT buffer for 10 min and incubated in NBT/BCIP (Roche) for the coloring reaction. Post fixation and PBST washes were done before imaging on agarose plates. *cFgf8* RNA probes were provided by Dr. John J. Young (Simmons University; Young et al., 2019).

### 3.6 Time-lapse imaging and wound healing assay

Whole mount cultures and live imaging were conducted as described in (Chapman et al., 2001; Atsuta et al., 2013). For time-lapse imaging of wound closure, a wound was mechanically created in the electroporated hindlimb ectoderm with a tungsten needle 30 min before the start of imaging. The embryos were collected using filter paper rings and placed on a 50 mm glass-bottom dish (MatTek Corporation). The ectoderm was imaged using a Spinning disk (Yokogawa) - confocal laser microscope (Nikon Ti2). Images were taken every minute with a Z-stack interval of 2 μm and a range of approximately 30 μm. Images were processed with NIS-Elements (Nikon) and Fiji software.

## ACKNOWLEDGEMENTS

We thank Drs. Michael Levin (Tufts University) and Daisuke Saito (Kyushu University) for helpful discussion, and MicRoN imaging core (Harvard Medical School) for assistance of live imaging. This work was supported by JSPS KAKENHI Grant Number JP20K22658 (to Y. A.), and NIH grant HD03443 (to C. J. T.).

---

**Supplemental movie 1**

*In vivo* time-lapse analysis using a cultured whole embryo, the limb periderm of which was labeled with Lifeact-mRuby. Frames were taken every minute with a 40x Plan Apochromat WI λS objective lens. Total movie length: 30min. Scale bar: 25 μm.

**Supplemental movie 2**

*In vivo* time-lapse analysis showing cellular behavior and actin dynamics of wound epithelial cells. F-actin is pseudo-colored by intensity of its signals. Frames were taken every minute with a 40x Plan Apochromat WI λS objective lens. Total movie length: 30 min. Scale bar: 25 μm.

**Supplemental movie 3**

*In vivo* time-lapse analysis reveals that epithelial cells at a wound edge seal a gap by extending actin-rich lamellipodia. F-actin is color-coded according to the intensity. Frames were taken every minute with a 40x Plan Apochromat WI λS objective lens. Total movie length: 90 min. Scale bar: 25 μm.

**Supplemental movie 4**

Misexpression of a dominant-negative form of Rac1 (dnRac1) inhibits the formation of lamellipodia during wound healing. F-actin is color-coded according to the intensity. Frames were taken every minute with a 40x Plan Apochromat WI λS objective lens. Total movie length: 90min. Scale bar: 25 μm.

